# A High-Confidence Atlas of Protein Methylation Enables AI-Driven Detection of Methylated Peptides

**DOI:** 10.64898/2026.07.01.733993

**Authors:** Shengbo Wang, Yannick Hartmaring, Christoph N Schlaffner, Emily H. Bowler-Barnett, Maria Martin, Jun Fan, Zhi Sun, Zhi Sun, Bernhard Y Renard, Eric W. Deutsch, Andrew R. Jones, Juan Antonio Vizcaíno

## Abstract

Lysine and arginine methylation regulate chromatin dynamics, transcription, and cellular signaling, however confident mass spectrometry (MS)-based detection and localization of this modification remain challenging. We reanalyzed eight public human methylation-enriched datasets using an open and standardized workflow that integrates database searching via the Trans-Proteomic Pipeline with a decoy-based statistical method for the independent estimation of false localization rates (FLR). This yielded a high-confidence Human Methylation Atlas of 1,828 sites (57 methyl-lysine, 1,771 methyl-arginine) across 1,021 proteins, classified into Gold, Silver, and Bronze confidence tiers. This is far fewer sites than reported in previous studies, reflecting the application of stringent FLR control, and what we hypothesise is potential high-false discovery in previous analyses.

We then leveraged this resource to adapt a deep learning-based methodology for the improved detection of methylated peptides. Three mouse methylation-enriched datasets were reanalysed to augment training, and the phosphoproteomics-trained AHLF (*ad hoc* learning of peptide fragmentation) model was fine-tuned by transfer learning to create AHLF-Methylation. The model achieved mean ROC-AUC values of 0.824 on human spectra, and 0.829 on combined human-mouse spectra. The atlas is available through PTMeXchange and PRIDE, with curated site evidence integrated into UniProt and PeptideAtlas.

## Introduction

Post-translational modifications (PTMs) play a central role in regulating protein function, stability, and interactions, thereby influencing nearly all aspects of cell biology [1, 2]. Among the diverse PTMs, methylation of lysine and arginine has been characterised as a critical modulator of epigenetic signaling, transcriptional regulation, and DNA damage response [3, 4], among other biological phenomena. The dysregulation of methylation pathways is a well-established driver of human diseases, most notably in cancer and neurodevelopmental disorders [5, 6]. This broad physiological and pathological significance has motivated efforts to comprehensively map methylation sites across the human proteome.

Mass spectrometry (MS)-based proteomics is the primary method for the large-scale study of PTMs [7]. However, the analysis of methylation presents distinct challenges. The detection and confident localization of methylated peptides is more complicated than other PTMs, not only by the mass shift itself (14.01565 Da), but also by the limitations of current enrichment strategies, as antibody-based enrichment methods often exhibit sequence biases and variable specificity, that can lead to incomplete identification of the methylated proteome [8]. Furthermore, methylated peptides often exhibit neutral loss of labile methyl groups upon collision-induced dissociation (CID), complicating confident site localization [9, 10]. While alternative fragmentation techniques like electron-transfer dissociation (ETD) can help mitigate this, the resulting data heterogeneity across existing protein methylation datasets is very high.

Public data repositories such as the PRIDE database [11] and other ProteomeXchange resources store large numbers of PTM-enriched datasets. Heterogeneous processing pipelines, different data formats, variable scoring thresholds, and the absence of widespread and harmonised PTM localization metrics (e.g., PTM site probability scores) make the comparison of the results included in different deposited studies impractical. To address this issue, consequently, reanalysis of public proteomics datasets is being increasingly used for data harmonisation purposes [12]. In the case of PTM-enriched datasets, data reanalysis can be performed to harmonize peptide/protein identification confidence, resolve ambiguous site localizations and correct false assignments arising from e.g. isobaric interferences or insufficient fragmentation [13]. Harmonised collections of PTM-enriched reanalysed datasets are being increasingly used for AI (Artificial Intelligence)-related applications [14–16].

In the context of the ‘PTMeXchange’ initiative we have already created a number of harmonised collections of reanalysed PTM-enriched datasets, including rice and *Plasmodium falciparum* phosphorylation [17, 18], human SUMOlyation [19] and human ubiquitination [14], among others. In all these studies, we used a statistical method based on the use of decoy modified amino acids for performing an independent estimation of false localization rate (FLR), providing a stringent, site-level confidence metric comparable across datasets [14, 18].

AI and deep learning (DL) in particular have emerged as powerful strategies for PTM identification by capturing complex spectral patterns that conventional algorithms often miss. More recently, Charih *et al.* introduced MethylSight 2.0, a sequence-based lysine methylation site predictor that leveraged protein-language-model embeddings, a transformer architecture, and multitask learning across lysine PTMs to achieve a significant leap in prediction accuracy of protein methylation, demonstrating the power of DL for this task [20]. While such computational predictors are invaluable for generating hypotheses, they ultimately require experimental validation. Furthermore, they cannot capture dynamic, context-specific methylation events that are revealed through direct proteomic measurements. A more direct strategy involves training models to interpret mass spectra. The *Ad hoc* Learning of Peptide Fragmentation (AHLF) framework exemplifies this approach, using a temporal convolutional network to learn diagnostic fragmentation patterns directly from spectra, enabling accurate detection of spectra coming from phosphorylated peptides [21]. We here propose adapting this powerful, spectrum-centric learning strategy to methylation proteomics.

A fundamental bottleneck, however, is the scarcity of high-confidence methylated-peptide spectra needed to train a robust model *de novo*. To overcome this, we employed a transfer learning strategy. We hypothesized that the foundational knowledge of general peptide fragmentation physics embedded within AHLF, pre-trained on massive phosphoproteomics datasets are transferable to protein methylation. By initializing our model with these learned features and fine-tuning it on our high-confidence methylation atlas, we can specialize it for methylation detection without requiring an impractically large training set. This approach directly addresses the data scarcity challenge and enhances the sensitivity of methylation site identification from MS data.

Here we have applied the PTMeXchange computation pipelines to reanalyze publicly available human and mouse methylation-enriched datasets. We provide: (i) a high-confidence atlas of human methylation sites; (ii) a demonstration of how this resource (including reanalysed mouse methylation datasets used solely for training) can empower DL through transfer learning, creating the model AHLF-Methylation, which is available to download in Github (https://github.com/shengbokw/AHLF-Methylation); and (iii) the public release of all data and models through PTMeXchange (https://www.proteomexchange.org/ptmexchange/index.html), PRIDE (dataset PXD076372), UniProt [22] and PeptideAtlas [23] to support future research.

## Materials and Methods

### Selection of datasets

Public datasets from the PRIDE database were selected for reanalysis based on the following criteria: (i) human/mouse-derived samples enriched for lysine or arginine methylation; (ii) Data Dependent Acquisition (DDA) data generated using Thermo Fisher Scientific instruments (this encompasses most DDA datasets, and has best compatibility with our analysis pipelines); and (iii) availability of experimental metadata, either through the original publication or by direct communication with the authors/data submitters.

Following manual curation, eight human methylation-enriched datasets were retained for reanalysis. The human datasets represent a range of biological contexts, including widely used cell lines (e.g., HEK293, HeLa, U2OS) and primary tissue samples. Where a dataset contained distinct experimental conditions (e.g., different treatments or types of enrichment), the raw files were processed separately to maintain condition-specific integrity. For instance, datasets PXD003700 and PXD027949 were each reanalyzed as three independent sub-experiments.

Additionally, three mouse (*Mus musculus*) methylation-enriched datasets were also reanalysed to augment the training data for DL purposes. A summary of the selected human and mouse datasets, including PRIDE accession numbers, key instrumental and biological characteristics is provided in Tables 1 and 2, respectively.

**Table 1.** List of human Methylation-enriched proteomics datasets reanalysed in this study and their main characteristics. A total of 1,099 raw files were reanalysed. The numbers of methylated sites, precursor mass tolerances, and fragment mass tolerances shown in the table are those reported in the original studies.

| Datasets | # of raw files | Methylated sites reported | Instrument | Quantification Label | Target | Peptide mass tolerance (ppm) | Fragment tolerance (Da) | Biological samples |
| --- | --- | --- | --- | --- | --- | --- | --- | --- |
| PXD003700 [24] | 140 | 7,382 | Q Exactive | SILAC | R | 7 | 0.02 | HEK293, HeLa, and U2OS cells |
| PXD005497 [25] | 241 | NA | Q Exactive | SILAC | K | 4 | 0.02 | K562 cells |
| PXD012007 [26] | 188 | NA | Q Exactive | SILAC | R | 7 | default | primary human AML samples |
| PXD022424 [27] | 75 | 1,031 | Orbitrap Fusion Lumos | SILAC | R | 20 | 0.02 | HEK293 cells |
| PXD023727 [28] | 11 | NA | Q Exactive | label free | K/R | 10 | 0.02 | HEK293T, HeLa, COLO320, Caco-2 and H4 cells |
| PXD027949 [13, 29, 30] | 416 | 1,198 | Q Exactive HF | SILAC | K/R | 6;6;20 | 0.02 | Hela, SK-OV-3, NB4, U2OS |
| PXD038203 [31] | 10 | 403 | Orbitrap Fusion | label free | R | 20 | default | human breast cancer tumors |
| PXD032819 [32] | 18 | NA | Q Exactive HF | label free | R | 10 | 0.02 | T-cell acute lymphoblastic leukemia |

**Table 2.** List of mouse methylation-enriched proteomics datasets reanalysed in this study and their main characteristics. A total of 42 raw files were reanalysed.

| Datasets | # of raw files | Methylated sites reported | Instruments | Quantification on Label | Target | Strain |
| --- | --- | --- | --- | --- | --- | --- |
| PXD028884 [33] | 12 | 1,150 | Q Exactive HF | Label free | K/R | CARM1 |
| PXD030423 [34] | 4 | NA | Orbitrap Fusion Lumos | Label free | R | KPC |
| PXD064423 [35] | 26 | NA | Orbitrap Fusion | TMT | R | Cas9-OT/P14+-I |

### Proteomics MS raw data processing

Raw mass spectrometry files (.raw) from each dataset were first converted to the mzML format using ThermoRawFileParser (version 1.3.4) [36]. To establish the appropriate search parameters, an initial subset of raw files from each dataset was processed using Fragpipe [37] in an open search mode to select the optimal search parameters. This step identified the potential modifications (as delta masses), to check the level of methylation enrichment of the datasets and determine the variable modifications to be included.

For the database search, all MS raw files were processed independently using the Comet search engine (version 2024) on a Linux-based high-performance computing cluster. Searches were performed against a combined database containing the UniProt human reference proteome (one canonical protein per gene, release 2024_01) or the UniProt mouse reference proteome (release 2025_04) and the cRAP sequences of common contaminants (https://www.thegpm.org/crap/, obtained April 2024). Decoy sequences for false discovery rate (FDR) estimation were generated by reversing the target sequences using the tool integrated within FragPipe.

Search parameters were explicitly configured to detect lysine (K) and arginine (R) methylation. We included the following variable modifications: mono- and di-methylation of lysine and arginine (+14.01565, +28.03130 Da). In addition to these, variable modifications identified in more than 1% of PSMs during the open search, along with oxidation of methionine and N-terminal protein acetylation, were included as search parameters in all searches. We added carbamidomethylation of cysteine as a fixed modification. The search was configured with trypsin as the protease, allowing for up to four missed cleavages. All other search parameters were set to Comet defaults. Post-processing and statistical validation (described in the following section) employed a decoy amino acid strategy using alanine (A) residues which was added on the Comet modification parameters for methylation, to independently estimate the false localization rate (FLR) for each identified methylated site [38]. This method provides a stringent, site-specific confidence metric by introducing decoy methylations on alanine, which is not a known target for biological methylation.

### Post-processing of the search results

Statistical validation of PSMs and distinct peptide sequences was conducted using PeptideProphet and iProphet from the Trans-Proteomic Pipeline (TPP version 7.1.0) [39]. High-confidence PSM matches were obtained, and PTM site localization was computed using PTMProphet (TPP), generating a unified mzIdentML format file [40] as the output.

The searching result files from the TPP were processed using a custom Python script (mzidFLR; https://github.com/PGB-LIV/mzidFLR), as previously described [38] and also applied in prior PTMeXchange projects (https://www.proteomexchange.org/ptmexchange/index.html). First, a global FDR was calculated at the PSM level at a 1% threshold to retain high-confidence matches. The results were converted to a site-based format, with PTM localization scores assigned to each methylation site. Contaminants, decoy hits, and non-methylated PSMs were excluded, retaining only methylated sites for downstream analysis. Second, to estimate the probability of correct PTM localization, the PTM localization probability (from PTMProphet) was multiplied by the PSM probability (from PeptideProphet). Redundancy in PSMs site-based evidence was addressed by collapsing the data to a peptidoform level using a binomial adjustment [17, 41]: the probability of a site being methylated was calculated by comparing the number of methylation events to the total evidence for that site across all PSMs. FLRs were estimated by introducing decoy alanine residues, yielding a 1% FDR PSM file and a PTM site-specific FLR file for quality control (QC) analysis for each individual dataset. To combine results across datasets, a meta-analysis approach was employed to control FLR inflation. PTM sites were classified into Gold, Silver, and Bronze categories. Classification thresholds were set as follows: Gold (threshold ≥ 2 datasets with FLR <1%), Silver (2 > threshold ≥ 1 dataset with FLR <1%) and Bronze (one or more datasets with FLR >1% and <5%).

### Label Generation for Training

#### Human Datasets

As a result from the reanalyses we had the FDR-filtered (<1% PSM-level) list of PSMs for each dataset, along with the high-confidence methylated-peptide identifications categorized into confidence tiers (Gold, Silver, Bronze). To create a high-quality dataset for model training, methylation assignments on C-terminal lysine or arginine peptide residues were discarded because tryptic cleavage at methylated Lys/Arg residues can occur but is less efficient than cleavage at unmodified residues, making C-terminal methyl-K/R assignments more difficult to interpret confidently [42]. PSMs from the Gold and Silver tiers were used to extract the corresponding experimental MS/MS spectra from the original MGF (Mascot Generic Format) files. Each spectrum was then labeled as “methylated” or “non-methylated” based on its associated PSM, whereas the “non-methylated” class comprised spectra from confidently identified, unmodified peptides within the same reanalyzed datasets. After filtering, a total of 58,486 spectra were obtained, the dataset was balanced between methylated and non-methylated spectra. To prevent peptide-sequence leakage between model-training and evaluation set, spectra were partitioned by peptide-sequence group rather than by individual spectrum. The spectra were then divided into training, validation and reserved test sets using an 80:10:10 split. This procedure ensured that no peptide sequence was shared between the training, validation, and the test sets, thereby guaranteeing a rigorous evaluation of the model’s capabilities.

#### Mouse Datasets

To expand the training dataset and evaluate the model’s ability to learn universal methylation fragmentation patterns across species, we integrated mass spectra from three mouse methylation-enriched datasets using the same processing workflow to ensure consistency. However, due to the small number of available mouse datasets and the lower abundance of high-quality methylation sites with FLR<0.01, we could not classify the mouse data into the Gold/Silver/Bronze categories used for human data. Instead, we employed a FLR-based thresholding strategy to generate reliable labels: spectra from methylation sites with FLR < 0.01 were labeled as true positives, while those with FLR > 0.38 were labeled as false positives (non-methylated), sites with intermediate FLR values (0.01 - 0.38) were excluded from training as ambiguous data. The 0.38 threshold was empirically determined to achieve a more balanced class distribution for model training. This conservative strategy yielded 1,324 true and 1,422 “false” mouse methylation PSMs, which were then merged with the human dataset for training purposes. The mouse data constituted approximately 5% of the human training set, which contained 29,246 true/false PSMs.

### Transfer Learning from the AHLF methodology

DL models for interpreting MS data rely on structured, efficient data representations. The AHLF architecture is designed to learn directly from MS/MS data [21]. First, mass spectra were transformed into a compact bi-vector representation. This method leverages the sparsity of centralized spectra by dividing the mass-to-charge (m/z) range (100–1,900 Da) into fixed 0.5 Da segments. Each segment retains the peak with the highest intensity, encoding its intensity alongside its precise position within the segment (m/z remainder) as a tuple. This ultimately generates a (3,600, 2) structured tensor for each spectrum, preserving the raw m/z values of selected peaks while being fully compatible with convolutional and recurrent neural network layers.

Training such a model from scratch requires hundreds of thousands of labeled data points, which do not currently exist for methylated peptides. Instead, we used transfer learning, initializing our model using the weights from the previously created AHLF models (AHLF-Phospho-α, -β, -γ, and -δ) [21]. These four variants of the model originated from the same architecture but were trained on different, non-overlapping splits of a large phosphopeptide dataset, providing diverse initializations for transfer learning. To ensure that the fine-tuned models captured methylation-specific rather than phosphorylation-specific spectral features, all variants were subsequently adapted exclusively using high-confidence methylated peptide spectra, as described below.

This foundation provided our model with a robust prior understanding of general peptide fragmentation patterns. We then fine-tuned this pre-trained model on our curated dataset of high-confidence methylated peptides mass spectra. To ensure stable and effective adaptation, we made three key hyperparameter adjustments: (i) Learning Rate: The learning rate was significantly reduced to facilitate gentle, incremental updates to the model weights, thereby preserving the foundational knowledge of peptide fragmentation learned from phosphoproteomics data while allowing for specialization to methylation-specific spectral features; (ii) Regularization: The dropout rate in the fully connected (dense) layers was increased to enhance regularization and mitigate the risk of overfitting on the smaller, methylation-specific training set; (iii) Layer Freezing: The initial feature-extraction layers (specifically the first four convolutional blocks) were frozen. This preserved the model’s ability to interpret low-level spectral features without altering the foundational weights, while the subsequent layers remained trainable to adapt to methylation-specific patterns. To determine the optimal fine□tuning hyperparameters, we performed a systematic grid search over three factors: learning rate (1e□4, 1e□5, 5e□5), random seed for training set shuffling (1, 42, 123), and the number of trainable dense layers (3 or 4) following the frozen convolutional blocks. All four AHLF□Phospho model variants (α, β, γ, δ) were fine□tuned for each combination, and performance was evaluated on the held□out test set using median ROC□AUC (Receiver Operating Characteristic - Area Under Curve). The combination of seed 42, a learning rate of 1e□4, and 5 trainable dense layers consistently yielded the highest median ROC□AUC across the four variants. These settings were therefore used for all subsequent fine□tuning experiments reported in this study.

The core transformer architecture and all other hyperparameters were kept consistent with the original AHLF implementation. Model performance was evaluated using standard metrics. The final, continuous prediction scores from each fine-tuned variant were converted to binary class labels using a discrimination threshold of 0.5, and the balanced accuracy, F1-score, and ROC-AUC were calculated using Scikit-learn [43].

## Results

### Systematic reanalysis reveals limited high-confidence methylation sites

We reanalyzed eight curated publicly available human methylation-enriched proteomics datasets (Table 1) using a standardized pipeline within the PTMeXchange framework (see ‘Material and Methods’) with unified FDR and FLR control. We identified a total of 1,921 high-confidence methylated peptides across all human datasets (summarized in Table 3). The subsequent application of a 1% FLR filter at the peptidoform-site level, a more rigorous metric for site-specific confidence, revealed significant variation in data quality across datasets. As detailed in Table 3, the number of confidently localized methylation sites at a 1% FLR level ranged from none (four datasets) to 928 (dataset PXD032819). Specifically, datasets PXD005497 and PXD023727 contained an insufficient number of sites passing the 1% FLR threshold and were therefore excluded from the final meta-analysis and categorization pipeline. This underscores the critical importance of reanalysis and unified statistical validation for data harmonisation.

**Table 3.** PSM counts for each human dataset at 1% FDR, methylated-peptide PSM counts at 1% FDR, site Counts (excluding pA decoy sites) for all PSMs collapsed by peptidoform position and at each of the FLR Thresholds: 1% and 5%.

| **Original dataset identifier** | **1% FDR PSM count** | **1% FDR Methylated-peptide count** | **Peptidoform-site count** | **1% FLR Peptidoform site count** | **5% FLR Peptidoform site count** |
| --- | --- | --- | --- | --- | --- |
| PXD003700 - GndCl | 184,948 | 38,740 | 11,542 | 612 | 1,191 |
| PXD003700 - RIPA | 166,540 | 39,918 | 11,512 | 56 | 2,550 |
| PXD003700 - siPRMT | 551,044 | 40,234 | 11,642 | 0 | 0 |
| PXD005497 | 1,424,643 | 9,835 | 3,655 | 0 | 0 |
| PXD012007 | 574,144 | 14,818 | 2,175 | 133 | 231 |
| PXD022424 | 1,094,885 | 124,049 | 6,965 | 869 | 3,033 |
| PXD023727 | 25,025 | 855 | 723 | 0 | 0 |
| PXD038203 | 57,608 | 798 | 401 | 3 | 3 |
| PXD032819 | 297,409 | 62,832 | 4,772 | 928 | 3,017 |
| PXD027949-Bremang | 1,785,320 | 43,363 | 13,981 | 18 | 19 |
| PXD027949-Spadotto | 242,694 | 1,231 | 659 | 0 | 0 |
| PXD027949-Massignani | 807,613 | 17,862 | 3,620 | 28 | 166 |

Methylation sites from the six remaining datasets (PXD003700, PXD012007, PXD022424, PXD027949, PXD038203, and PXD032819) were classified into a three-tier confidence system (Gold, Silver, Bronze, see ‘Material and Methods’). The Gold set (sites identified in at least 2 datasets at <1% FLR) contained 156 high-confidence sites. The Silver set (sites in one dataset at <1% FLR) contained 362 sites, and the Bronze set (sites at <5% FLR) contained 1,310 sites (Fig. 1A).

**Figure 1.**
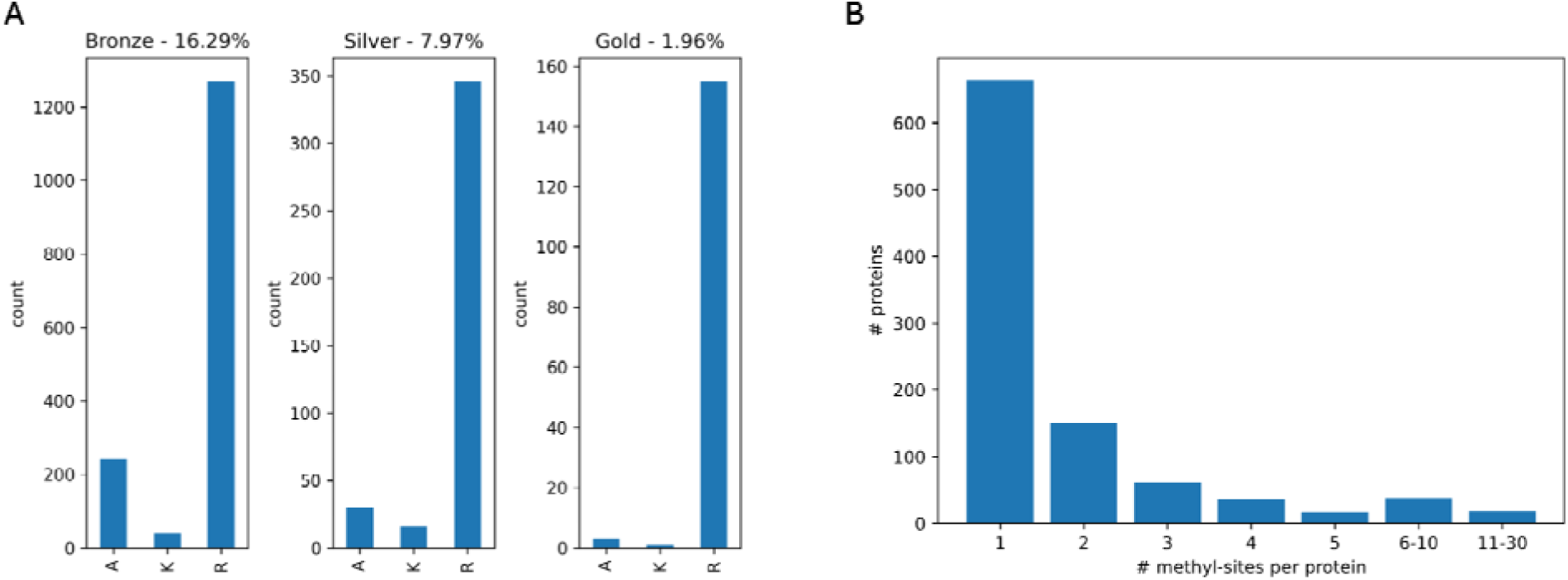
Characterization of the high-confidence human methylation atlas. **A)** Distribution of high□confidence methylated protein sites across confidence tiers (Gold, Silver, Bronze) and residue types (Arginine/R, Lysine/K, Alanine/A), highlighting the predominance of arginine methylation (datasets PXD003700, PXD012007, PXD022424, PXD027949, PXD038203, PXD032819). **B)** Number of identified methylation-sites per protein.

Notably, arginine methylation (R) sites were more prevalent and reproducible in the combined results than lysine (K) methylation sites, with 155 Gold-tier R sites compared to a single Gold-tier K site (Fig. 1A). This likely reflects the biological context of the selected studies (enriched for arginine methylation via PRMT inhibition). A small number of sites were localized to the decoy amino acid alanine (A), which are considered false positives and were excluded from the downstream analysis. The distribution of methylation sites across proteins (Fig. 1B) reveals that while most methylated proteins contain a single modified residue, a subset is heavily methylated.

To enable cross-species comparisons and augment our training data, we applied the same reanalysis pipeline to three mouse methylation-enriched datasets (PXD028884, PXD030423, PXD064423). The results of the reanalysis for each dataset are summarised in Table 4.

**Table 4.** PSM counts for each mouse dataset at 1% FDR, methylated-peptide PSM counts at 1% FDR, site counts (excluding pAla decoy sites) for all PSMs collapsed by peptidoform position and at each of the FLR thresholds (1% and 5%).

| Original data set identifier | 1% FDR PSM count | 1% FDR Methylated-peptide count | Peptidoform-site count | 1% FLR Peptidoform site count | 5% FLR Peptidoform site count |
| --- | --- | --- | --- | --- | --- |
| PXD028884 | 96,749 | 9,174 | 1,199 | 27 | 56 |
| PXD030423 | 4,936 | 434 | 187 | 19 | 19 |
| PXD064423 | 59,347 | 8,254 | 255 | 3 | 3 |

### Transfer Learning Enables Effective Detection of Methylated Peptides

The limited scale of high-confidence mass spectra coming from methylated peptides presents a challenge for training DL models *de novo*. The AHLF model was originally developed and trained on a massive corpus of phosphorylated peptide mass spectra [21], learning to recognize complex fragmentation patterns that are fundamental to peptide behavior in the mass spectrometer. Given that some of the foundational fragmentation principles are shared across PTMs, we hypothesized that this pre-trained model could be adapted for methylation detection using a transfer learning approach. We fine-tuned the AHLF model—initially pre-trained on a vast corpus of phosphopeptide spectra—using our curated dataset of 58,486 high-confidence methylated-peptide spectra to create the model AHLF-Methylation.

To robustly evaluate performance, we fine-tuned four pretrained AHLF variants (AHLF-Phospho-α, -β, -γ, and -δ) on the curated methylation dataset, yielding four independently adapted AHLF-Methylation models. The methylation spectra were split into training, validation, and reserved test sets using peptide-sequence-group partitioning, ensuring that spectra assigned to the same peptide sequence group were not shared across splits. This design reduced the risk of peptide-sequence leakage between training and evaluation sets.

Hyperparameter optimization (learning rate, random seed, and number of trainable dense layers) identified seed 42, a learning rate of 1e□4, and 4 trainable dense layers as the best combination (see Supplementary File 1). These settings were used for all models reported below. The results are summarized in Table 5.

**Table 5.** Performance metrics of the 4 fine-tuned AHLF-Methylation models on the held-out test set of mass spectra from methylated peptides.

|  | ROC-AUC | Accuracy | F1 Score | Precision | Recall | Bacc |
| --- | --- | --- | --- | --- | --- | --- |
| Alpha model | 0.834 | 0.732 | 0.706 | 0.782 | 0.643 | 0.732 |
| Beta model | 0.852 | 0.745 | 0.722 | 0.792 | 0.663 | 0.745 |
| Delta model | 0.791 | 0.698 | 0.672 | 0.735 | 0.619 | 0.698 |
| Gamma model | 0.815 | 0.686 | 0.625 | 0.774 | 0.524 | 0.686 |

The fine-tuned model demonstrated a significant enhancement in performance for detecting methylated peptides compared to its base state. AHLF-Methylation achieved a median ROC-AUC of 0.824 and a median F1-score of 0.689 on a held-out test set (Supplementary Information Section 1 and Table S1). This performance, achieved with a relatively small training set (6 datasets), highlights the efficacy of transfer learning for across spectra coming from different PTMs.

A direct comparison of AHLF-Methylation with existing computational tools for protein methylation detection is not feasible since, to the best of our knowledge, no currently available method addresses the identical task of classifying whether a given mass spectrum originates from a methylated peptide. The most closely related approaches, such as MethylSight [44], operate at the sequence level by predicting methylation-prone residues within protein sequences, and therefore address a fundamentally different problem.

To further evaluate the data efficiency of our approach, we reduced the size of the methylated-peptide training set by different percentages rather than by simply varying the number of constituent datasets and assessed performance across training fractions from 100% to 10%. Four pretrained AHLF variants (α, -β, -γ, and -δ) were fine-tuned on each reduced subset and evaluated using the same validation framework. Performance was summarized across models using both mean and median ROC-AUC values (Supplementary File 2; Figure 2). The full training set yielded the highest mean ROC-AUC (0.8229), while reduced subsets generally showed lower performance, 90% (0.8153), 80% (0.8113), 70% (0.8100). Performance declined further at 60% (0.7880), but remained comparatively higher at 50% (0.7792), 40% (0.7729), 30% (0.7713), and 10% (0.7768), whereas the lowest mean ROC-AUC was observed at 20% (0.7332). Median ROC-AUC values showed a similar pattern, with 90% (0.8200) and 100% (0.8243) performing best overall.

**Figure 2.**
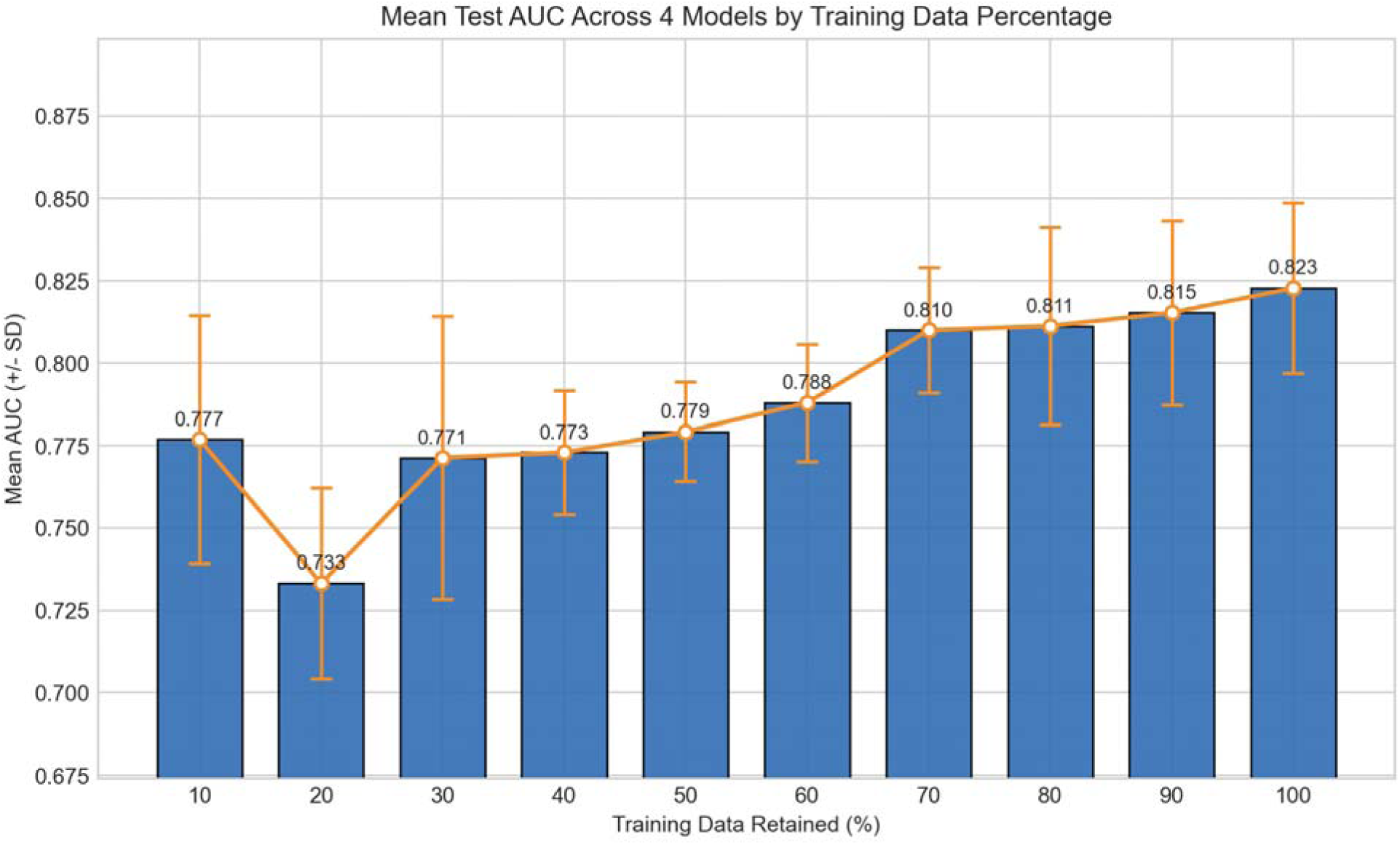
Mean test AUC across four fine-tuned models as a function of retained training data. Error bars denote the standard deviation across the four model variants.

This pattern indicates that reducing the amount of training data generally lowers performance, although the relationship is not strictly linear. The 10% subset behaved as an outlier, yielding stronger performance than the 20% setting and some intermediate fractions. This suggests that, in addition to training-set size, the exact peptide composition retained in each subset exerts a substantial influence on downstream performance. Run-to-run and model-to-model variability are also likely to contribute to the observed fluctuations. Therefore, these results should not be interpreted as evidence that smaller training sets are inherently superior to the full dataset, but rather as an indication that fine-tuning performance is shaped by both data quantity and subset composition. Together, these findings emphasize that, in PTM-specific transfer learning, data quality, harmonized curation, and balanced sampling strategies may be as important as the overall training-set size.

### Model interpretation indicates preferential use of peptide-informative fragment ions

To assess whether AHLF-Methylation predictions were supported by peptide-informative spectral evidence, we performed a model-interpretation analysis following the strategy used in the original AHLF-Phosphorylation study [21]. For each spectrum, peak-level SHAP values were computed and mapped back to experimental spectral features. Peaks with high model attribution were then compared with theoretical fragment ions expected from the database-assigned peptidoform. This enabled us to evaluate whether model-emphasised peaks were enriched among peptide-derived fragment ions rather than being dominated by unexplained spectral features.

To quantify this relationship, we calculated two complementary spectrum-level ratios. The SHAP-value ratio was defined as the proportion of absolute SHAP attribution from the intensity channel that was assigned to peaks matching theoretical fragment ions. The intensity ratio was defined as the proportion of total measured spectral intensity contributed by those same matched peaks. Thus, the SHAP-value ratio reflects the extent to which the model assigns importance to peptide-related peaks, whereas the intensity ratio reflects the abundance of those peaks in the observed spectrum.

Across the evaluated spectra, the overall intensity-channel SHAP-value ratio was modestly higher than the corresponding intensity ratio (Supplementary File 3). This indicates that the model AHLF-Methylation did not simply prioritize the most intense peaks in each spectrum. Instead, peaks consistent with theoretical peptide fragment ions contributed disproportionately to the prediction signal, even when those peaks did not dominate total spectral intensity. This supports the view that the fine-tuned classifier is using chemically meaningful spectral evidence. However, this analysis does not by itself reveal which exact fragmentation relationships, ion series, or methylation-dependent peak shifts are most important for classification.

### Incorporation of Cross-Species Mouse Data Modestly Augments Model Performance

To validate whether integrating evolutionarily conserved methylation patterns could enhance model robustness, we expanded the training set by adding high-confidence mouse methylated-peptide spectra to the human ones (see ‘Material and Methods’). After preprocessing, the mouse datasets contributed 1,324 true and 1,422 false methylated-peptide mass spectra, which was used to fine-tune the four AHLF model variants (AHLFp-α to δ) by merging into the human dataset, producing the AHLF-Methyl-HM models. The performance of the resulting AHLF-Methyl-HM models is summarized in Supplementary Information Section 2 and Table S2.

Across the four human–mouse fine-tuned models, performance remained robust, with a mean ROC-AUC of 0.829, mean F1 score of 0.693, and mean balanced accuracy of 0.725. The Beta variant achieved the highest ROC-AUC (0.851), indicating the strongest overall discrimination between methylated and non-methylated spectra. At the fixed classification threshold, the Alpha variant showed the strongest classification performance, with the highest accuracy (0.753), F1 score (0.736), recall (0.694), and balanced accuracy (0.753). The Delta and Gamma variants showed comparatively weaker performance, with Delta yielding the lowest ROC-AUC (0.801) and Gamma showing the lowest recall (0.524) and balanced accuracy (0.685). Taken together, these results indicate that mouse methylated-peptide spectra can be incorporated into the training workflow without substantially compromising predictive performance. The modest differences between model variants suggest that the effect of cross-species data integration depends on the specific pretrained initialization and fine-tuning trajectory. Overall, these findings support the use of carefully curated model-organism datasets as supplementary training data for spectrum-based peptide methylation detection.

## Discussion

In this study we systematically reanalyzed publicly available MS-based proteomics datasets to construct a highly reliable human protein methylation atlas. We then used the results to improve the detectability of methylated-peptide mass spectra, using a transfer-learning-based approach.

This study highlights the critical role of systematic reanalysis in maximizing the value of public proteomics data. Two of the eight human datasets were excluded from the final results due to insufficient site counts after filtering through a stringent 1% FLR [17]. Similarly, the three mouse datasets, while contributing valuable spectra, yielded relatively few sites passing the 1% FLR threshold (ranging from 3 to 27 sites per dataset). This threshold does not constitute a comprehensive assessment of data quality but is a necessary step to ensure downstream interpretation is grounded in a reliable localization of PTMs, in this case methylation. As in previous studies [17, 18], our FLR estimation method provides transparent and comparable site confidence metrics, effectively mitigating the risk of false positives.

However, in the case of methylation, the proportion of sites that remained after the application of the decoy-aminoacid methodology is considerably lower than for other PTMs (considering phosphorylation, ubiquitination and SUMOylation), as found in previous analogous studies using the same methodology [14, 19, 41]. We speculate that the main reason is that the FLR is indeed high, due to all the possible ways in which a mass shift could get confused by a search engine (Supplementary Information Section 2 and Table S3). Methylation-like mass shifts can also arise from alternative peptide interpretations, particularly when homologous or isoform-derived peptides differ by amino acid substitutions with similar mass differences (Table S3). Potential errors can then arise from e.g. expanded variable-modification search spaces, incorrect modification localization, homologous peptide sequences, and artefactual rather than biological modifications. The 14 Da mass shift is indeed much smaller than that introduced by other PTMs such as phosphorylation, ubiquitination (GlyGly) or SUMOylation remnants.

It would also be possible that the decoy-amino acid method is not as well suited for methylation as it is the case for other PTMs, although we do not believe this to be likely. As indicated above, the method has been successfully applied for phosphorylation, as well as other lysine modifications ubiquitination and SUMOylation - leading to large and robust collections of high-quality PTM builds being created. A more likely explanation in our view is that a standard search for methylated peptides has very high FDR, far exceeding the 1% FDR estimated by the target-decoy method as noted above and in a previous study [45]. The use of the decoy amino acid then manages to control for this, since in false positive peptide identifications, site localisation is often confidently assigned to the best fitting amino acid i.e. A (decoy), K or R (targets) at random - leading to exceptionally high FLR estimates. The decoy-amino-acid FLR strategy is therefore expected to remove not only low-quality spectra, but also high-scoring methylation-like assignments with plausible competing explanations. This supports the conservative nature of the final high-confidence methylation atlas.

Although the primary aim of this study was to generate a high-confidence, AI-ready human methylation build, a second contribution of this study is the demonstration that transfer learning can be used to develop a methylation-specific spectral classifier despite the limited availability of high-confidence methylated peptide spectra. By fine-tuning AHLF-Phospho on our methylation atlas, we generated the model AHLF-Methylation, showing that spectral representations learned in one PTM-related domain can be adapted to another. This is consistent with the original AHLF framework, which showed that a phosphopeptide-trained model could be transferred to the distinct task of cross-linked peptide detection [21], as well as with our additional recent application of the same framework to ubiquitination and acetylation [46]. In practical terms, AHLF-Methylation is best viewed as a spectrum-level prioritization tool that can pre-score tandem mass spectra according to their likelihood of representing methylated peptides, thereby enriching candidate spectra for downstream identification, targeted inspection, or follow-up analysis. More broadly, these findings suggest that the generalized peptide fragmentation representations learned by AHLF may be transferable across diverse analytical contexts. Our results extend this principle to methylation by taking advantage of shared fragmentation characteristics across PTMs, allowing the model to first learn general spectral features from the more abundant phosphorylation data before being specifically optimized for methylation. Although further evaluation across additional datasets and modification types will be important, the present results support transfer learning as a promising strategy for improving detection of understudied PTMs when modification-specific training data are limited.

Additionally, the SHAP-based interpretation analysis provides an important sanity check for AHLF-Methylation. The observation that model attribution was enriched among peaks matching theoretical peptide fragment ions suggests that the fine-tuned model relies, at least in part, on chemically meaningful spectral evidence rather than primarily on unexplained intensity patterns or dataset-specific artefacts. However, this analysis does not by itself establish that the model has learned methylation-site-specific diagnostic ions; rather, it supports the suitability of the mass spectra from methylated peptides as an AI-ready resource for spectrum-based methylated peptide detection.

Several limitations in our study must be acknowledged. First, the scale and diversity of the underlying datasets influence the conclusions. The absolute predominance of arginine methylation sites in the ‘Gold’ set reflects the biological research focus of existing public datasets (e.g., due to PRMT inhibition studies). Second, it is worth highlighting the imbalance of the available training data. Compared to phosphorylation (the data used to train the original AHLF-Phospho model), the number of confidently localized spectra from methylated peptides is small, leading to a strong bias toward negative (unmodified) labels. Such imbalance can hinder DL models, which tend to favor the majority class, reducing sensitivity for detecting true spectra coming from methylated peptides. The employed transfer learning strategy partially mitigates this issue by leveraging pretrained spectral representations and requiring fewer methylated examples to achieve robust classification. Nonetheless, the scarcity of positive labels (i.e. spectra from methylated peptides) remains a bottleneck, and future efforts should explore methods such as data augmentation, synthetic spectrum generation, and cost-sensitive learning to further counteract class imbalance.

Our exploratory integration of additional peptide methylation data coming from mouse highlights both the potential of cross-species training in DL approaches and the subtle challenges involved. The successful maintenance of high performance in the Beta variant (AHLF-Methylation-HM-β) suggests that the core fragmented patterns learned by the model are evolutionarily conserved between humans and mice, enabling effective knowledge transfer. This finding supports the biological principle that fundamental peptide chemical properties and enzyme recognition motifs can be conserved across species. However, the performance differences across model architectures, particularly the significant decline in recall for the Delta variant of the model, indicates that not all learned representations can generalize equivalently. The relatively limited mouse data (approximately 5% of the total training set) may also constrain its ability to significantly alter model decision boundaries, explaining why overall performance remained stable without achieving breakthrough improvements.

This work opens multiple avenues for future research. In the context of the PTMeXchange project, we will continue to create high-quality collections of PTM data thereby establishing a comprehensive AI-ready PTM resource. As more methylation-enriched datasets emerge from diverse disease states and cellular environments, integrating this context-specific information is essential for transitioning from static mapping to a dynamic understanding of methylation signaling in health and disease. Furthermore, we expect that integrating these high-confidence PTM maps will advance systematic investigations into PTM cross-regulation—a critical cellular regulatory layer that remains challenging to study robustly.

## Supporting information

Supplementary File 1 reports the hyperparameter-screening results used for AHLF-Methylation fine-tuning.

Supplementary File 2 reports the per-model results of the training-data reduction experiment.

Supplementary File 3 summarizes the SHAP-based model interpretation analysis.

## Acknowledgements

We would like to thank all data submitters who made their datasets publicly available through PRIDE and ProteomeXchange. J.A.V. and S.W. would like to acknowledge funding from BBSRC [grant numbers BB/S01781X/1, BB/Y513829/1], EPSRC [grant numbers EP/Y035984/1], Wellcome [223745/Z/21/Z] and EMBL core funding. A.R.J. would like to acknowledge funding from BBSRC [BB/S017054/1]. E.W.D. and Z.S. acknowledge National Science Foundation grant DBI-2324882. This work is also supported by a European Research Council (ERC) grant (eXplAInProt, grant number 101124385) to B.Y.R.

