## Supplementary Information for "A High-Confidence Atlas of Protein Methylation Enables AI-Driven Detection of Methylated Peptides"

**Authors and Affiliations**

^1^ European Molecular Biology Laboratory - European Bioinformatics Institute (EMBL-EBI), Wellcome Genome Campus, Hinxton, Cambridge, CB10 1SD. United Kingdom.

^2^ Hasso Plattner Institute for Digital Engineering, Digital Engineering Faculty, University of Potsdam, Potsdam, Germany.

^3^ Institute for Systems Biology, Seattle, Washington 98109, United States.

^4^ Institute of Systems, Molecular and Integrative Biology, University of Liverpool, Liverpool L69 7ZB, United Kingdom.

*Corresponding authors: Prof. Andrew R. Jones and Dr. Juan Antonio Vizcaíno

Contents

**Appendix Table S1**: Summary of key performance metrics for AHLF-Methylation and AHLF-Phospho models**.**.	2

**Appendix Table S2**: Performance of AHLF-Methyl-HM models fine-tuned on combined human and mouse methylation datasets……………………………………………………………...…….3

**Appendix Table S3**: Table showing potential alternative explanations for apparent +14.01565 Da methylation-like mass shifts	5-6

### **1. Key performance metrics for the AHLF-Methylation Models**

Supplementary Table S1 summarizes the median performance metrics of the fine-tuned AHLF-Methylation models in comparison with the original AHLF-Phospho models. The AHLF-Methylation models achieved a median ROC-AUC of 0.824, median F1 score of 0.689, and median balanced accuracy of 0.710. As expected, these values were lower than those of the original AHLF-Phospho models, which were trained and evaluated in the context of substantially larger phosphoproteomics datasets. This comparison is therefore intended to provide a reference point for model adaptation rather than a direct task-equivalent benchmark.

|  | **Median ROC-AUC** | **Median F1 score** | **Median Bacc** |
| --- | --- | --- | --- |
| AHLF-Methylation | 82.43 | 68.90 | 71.0 |
| AHLF-Phospho | 88.48 | 83.32 | 76.80 |

**Table S1. Summary of key performance metrics for AHLF-Methylation and AHLF-Phospho models.** Median ROC-AUC, F1 score, and balanced accuracy are shown for the fine-tuned AHLF-Methylation models and the original AHLF-Phospho models. AHLF-Phospho is included as a reference for the pretrained model family, while AHLF-Methylation reflects adaptation to the smaller methylation-specific training set.

Supplementary Table S2 reports the performance of the four AHLF-Methyl-HM models fine-tuned using the combined human and mouse methylated-peptide spectra. Across the four model variants, predictive performance remained stable, with mean accuracy of 0.726, mean F1 score of 0.693, mean balanced accuracy of 0.725, and mean ROC-AUC of 0.829. The Beta variant achieved the highest ROC-AUC, whereas the Alpha variant showed the strongest accuracy, F1 score, recall, and balanced accuracy at the selected classification threshold. These results indicate that incorporation of mouse methylation-enriched spectra did not substantially compromise model performance and may provide useful supplementary training data for spectrum-based methylation detection.

| **Human + Mouse** | **Accuracy** | **F1 Score** | **Precision** | **Recall** | **Bacc** | **ROC-AUC** |
| --- | --- | --- | --- | --- | --- | --- |
| Alpha model | 0.7534 | 0.7357 | 0.7834 | 0.6944 | 0.7528 | 0.8424 |
| Beta model | 0.7506 | 0.7292 | 0.7884 | 0.6783 | 0.75 | 0.8511 |
| Delta model | 0.7121 | 0.6838 | 0.7493 | 0.6288 | 0.7113 | 0.8006 |
| Gamma model | 0.6868 | 0.6234 | 0.7697 | 0.5239 | 0.6851 | 0.8218 |

**Table S2. Performance of AHLF-Methyl-HM models fine-tuned on combined human and mouse methylation datasets.** Accuracy, F1 score, precision, recall, balanced accuracy, and ROC-AUC are reported for the four AHLF model variants after fine-tuning on the combined human–mouse dataset.

### **2. Possible alternative explanations for a +14 Da mass shift**

Mass shifts of +14.01565 Da are commonly interpreted as mono-methylation of lysine or arginine residues. However, methylation-like mass shifts can also arise from alternative peptide interpretations, particularly when homologous or isoform-derived peptides differ by amino acid substitutions with similar mass differences (Supplementary Table S3). This issue is especially relevant for methylation proteomics because previous work has shown that methylated-peptide identifications can have substantially higher false discovery rates than implied by global PSM-level FDR estimates [1]. Such errors can arise from e.g. expanded variable-modification search spaces, incorrect modification localization, homologous peptide sequences, alternative arrangements of modifications, and artefactual rather than biological modifications.

To explore the potential scale of this ambiguity, we performed an exploratory *in silico* analysis of tryptic peptide pairs in a UniProt 100K proteins Swiss-Prot, TrEMBL, and isoform-containing search set. We identified 5,885 peptide pairs differing by a single amino acid substitution with a mass difference matching the mono-methylation mass shift of 14.01565 Da. When allowing for apparent mass differences compatible with +1 or +2 precursor isotope-selection errors, this increased to 23,536 peptide pairs. These theoretical pairs represent cases in which a lighter peptide plus an assigned +14 Da modification could potentially compete with an unmodified heavier peptide during database searching. Such ambiguity may be particularly problematic if the correct sequence is absent from the search database, for example due to missing protein isoforms/variant sequences, if the precursor isotope assignment is incorrect, or if fragment-ion evidence is insufficient to distinguish between closely related sequence alternatives.

A representative example is provided by the HMGB (High Mobility Group Box)-family peptides, where the peptides MSAYAFFVQTCR and MSSYAFFVQTCR occur in closely related protein sequences. In this case, an assignment such as MSA[Methyl]YAFFVQTCR can approximate the mass of the unmodified serine-containing peptide (MSSYAFFVQTCR) when isotope-error tolerance is allowed. Because these candidate peptides share most of their C-terminal sequence, many dominant fragment ions may be common between the two interpretations, while the sequence-discriminating N-terminal ions may be weak or absent. As a result, the spectrum can appear to be a high-quality PSM while remaining ambiguous with respect to the true peptide sequence and methylation localization.

Additional sources of apparent +14 Da shifts include sample-preparation artefacts such as methyl esterification of acidic residues, cysteine alkylation-related artefacts, incorrect modification identity, and incorrect localization of a true modification within the peptide. Several of these ambiguity classes have been described previously in large-scale methylation analyses, including isobaric peptide alternatives, incorrect methylation-site localization, cysteinyl-S-β-propionamide-containing peptides, and methylated acidic residues [1].

These observations provide a plausible explanation for the high FLR (False Localisation Rates) estimates observed for some methylation-like assignments. If a +14 Da mass shift can be explained by multiple sequence or artefactual alternatives, the search engine may assign the modification to lysine, arginine, or decoy residues depending on small differences in scoring and fragment-ion support. The decoy-amino-acid FLR strategy that we have used in this study is therefore expected to remove not only low-quality spectra, but also high-scoring methylation-like assignments with plausible competing explanations. This supports the conservative nature of the final high-confidence methylation atlas and helps explain why the number of retained sites is lower than in less-stringently filtered methylation resources.

| **(Alternative) explanation** | **Mass-shift logic** | **Consequence for methylation assignment** |
| --- | --- | --- |
| True Lys/Arg methylation | Addition of CH₂-like methyl group, +14.01565 Da | Correct biological interpretation if supported by site-determining ions |
| Homologous or isoform peptide differing by one CH₂ unit | Common residue substitutions such as Gly→Ala, Ser→Thr, Asp→Glu, Asn→Gln, or Val→Leu/Ile differ by ~14.01565 Da | A lighter peptide plus methylation can mimic an unmodified heavier peptide |
| Precursor isotope-selection error | Incorrect monoisotopic precursor assignment can shift the apparent precursor mass by one or more isotope spacings | Near-isobaric peptide alternatives may be brought within precursor tolerance |
| Decoy A[Methyl] versus Ser-containing peptide | Ala[Methyl] differs from Ser by ~1.98 Da, close to two isotope spacings | A two-isotope precursor error can make an unmodified Ser peptide resemble an A[Methyl] assignment |
| Methyl esterification of Asp/Glu | Sample-preparation artefacts can introduce +14 Da methyl ester modifications on acidic residues | Artefactual methylation can be misinterpreted as biological methylation if localized incorrectly |
| Cys alkylation artefacts | Propionamide on Cys is +14.01565 Da relative to carbamidomethylated Cys | A Cys artefact can mimic nearby Lys/Arg methylation |

**Table S3.** Potential alternative explanations for apparent +14.01565 Da methylation-like mass shifts. The table summarizes known ambiguity classes described in previous methylation-proteomics studies, standard mass spectrometry-based protein modification annotations, and the exploratory *in silico* peptide-pair analysis performed in this study.
